# A novel phylogenetic comparative model reveals strong signals for early high disparity in ichthyosaurs

**DOI:** 10.64898/2026.09.25.753454

**Authors:** Ricardo C. Ely, Fabio A. Machado

## Abstract

Early high disparity, where clades reach maximum morphological diversity early in their history, is often observed in the fossil record. Since there are currently no suitable methods to model these Early High Disparity (EHD) processes accurately, we introduce a new phylogenetic comparative method (PCM) to address this gap by combining two evolutionary modes—Ornstein-Uhlenbeck (OU) and Early Burst (EB)—leading to an initial surge in disparity until an early maximum, driven by the EB component, after which disparity declines due to OU constraints. We tested the EHD model on ichthyosaur skull length data and found it to be the best fit in nearly all cases compared to other modes. The median time of peak disparity (T_EHD_) is about 231.1 million years ago, during the Triassic, roughly 12.4% into the ichthyosaur lineage. To explore how environmental and geological factors might influence peak disparity, we performed changepoint analyses on sea level, sea surface temperature, rock area, and sedimentary sequence ages. While changepoints in temperature at 237.2 Ma and sequence boundaries at 238 Ma were closest to T_EHD_, no significant changepoints appeared to match T_EHD_.

**Significance Statement:** Early phylogenetic comparative methods used evolutionary models where trait variance (disparity) either gradually increased or remained stable around certain values. These models are still widely employed today, mainly with datasets of living traits. However, the fossil record shows many cases where clades do not follow the assumptions of continuous disparity increase or stability. For example, some fossil clades exhibit high disparity early on, then decline. We present a new PCM that specifically models such Early High Disparity, broadening the range of available models and allowing fitting of models supported exclusively by fossil data.

## Introduction

Ichthyosaurs were among the most successful and long-lived marine vertebrate radiations in Earth’s history, originating within ∼2 million years of the Triassic Period in the aftermath of the End-Permian Mass Extinction (EPME) (1). Even more striking than their rapid appearance was the extraordinary morphological disparity they achieved early in their history (2– 4). Early Triassic ichthyosauriforms ranged from the small-bodied, filter-feeding Hupehsuchus with cetacean-like adaptations (5), eel-like and lizard-like ichthyosaurs (6), to streamlined pelagic morphologies resembling later Jurassic and Cretaceous ichthyosaurs (1, 7). They also spanned an exceptional range of body sizes, from small taxa such as *Cartorhynchus* (8), *Chaohusaurus* (9), and *Grippia* (10), to the ∼17-meter-long Cymbospondylus youngorum [247.2-237 Ma (4)].

This rapid expansion in form and size indicates an intense adaptive radiation following the EPME. Ichthyosaurs reached peak body-size disparity within the first ∼10 million years of the Triassic, far earlier than cetaceans, which achieved their greatest size disparity over ∼53 million years from their Eocene origins to the present (4). This early burst produced high morphological disparity among basal ichthyosaurs, followed by a subsequent decline. Possible drivers of this decline include competition with newly diversifying marine reptiles (8–11), late Triassic sea-level regression that may have eliminated nearshore taxa while favoring pelagic forms (12), the end-Triassic mass extinction, which sharply reduced disparity and prevented ichthyosaurs from regaining their Triassic peak during the rest of the Mesozoic (2).

Such ‘early high disparity’ is frequently encountered across fossil clades (13–17). The fossil record reveals disparity peaks relatively early in the history of several clades; classic cases include the whole of arthropoda (13), Paleozoic rostroconchs (18), Paleozoic gastropods (14), crinoids (19), and Paleozoic blastozoans (20). Since the suggestion of a substantial early peak in disparity in Cambrian relative to Recent arthropods (13), the bulk of disparity studies have focused on fossil invertebrates. Vertebrates displaying early high disparity include carnivorans (21), possibly more specifically borophagines (16), ichthyosaurs, certain ceratopsian dinosaur subclades, stegosaurians, certain temnospondyl amphibian clades, among others (16, 17).

Several mechanisms have been proposed to explain this pattern, including post-extinction adaptive radiations(22, 23), developmental constraints (14), and selective extinctions (24, 25). Yet despite the apparent recurrence of early high disparity across the fossil record, existing phylogenetic comparative models only capture components of the pattern, such as early bursts of evolution (26, 27). No method has yet been proposed to explicitly model the full dynamic of an early rise to peak disparity followed by a subsequent decline.

Here, we introduce a new phylogenetic comparative method designed to model this dynamic explicitly, using ichthyosaurs as a focal test case. The model combines two established stochastic processes: the Ornstein-Uhlenbeck (OU) model (28, 29), commonly used to represent stabilizing selection toward an adaptive optimum (46, 47, 48, 50–52) and the Early Burst (EB) model, which is often used to describe adaptive radiations in which trait variance expands rapidly early in a clade’s history before slowing through time (27, 30). By combining these processes, the model captures an initial EB-like phase of rapid trait diversification, followed by a decline in disparity as OU-like constraints increasingly dominate (Fig 1). We refer to this combined process as the Early High Disparity (EHD) model.

**Figure 1.**
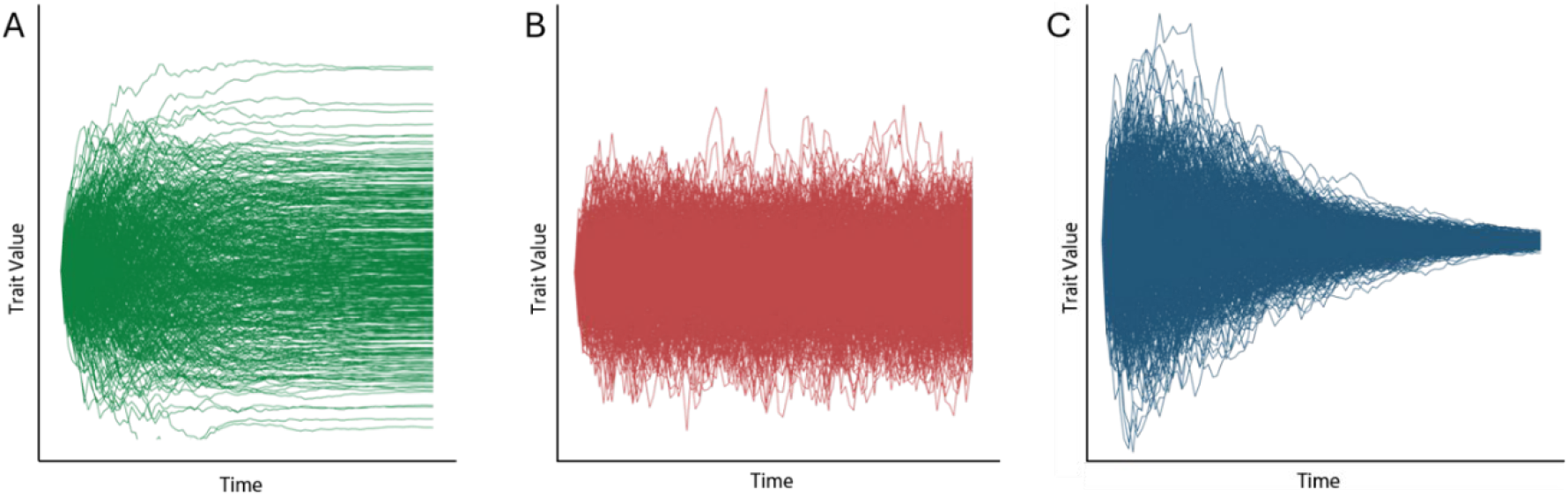
Monte Carlo simulations of A.) EB, B.) OU, and C.) EHD processes. Each set of Monte Carlo simulations represents 1000 time-series of ‘random’ walks, run for 100 time-steps.

Ichthyosaurs provide a strong empirical system for testing this model because they have repeatedly been described as exhibiting early peaks in disparity (3, 34, 53). Several studies suggest that ichthyosaurs reached peak disparity relatively soon after their origin (3, 53), potentially reflecting an EHD mode of evolution (34). Here, we explicitly test that hypothesis by applying the newly proposed EHD model to ichthyosaur trait evolution in a phylogenetic comparative framework.

## Results

We first evaluated whether ichthyosaur skull-length evolution is best described by an Early High Disparity (EHD) pattern. A phylomorphospace-through-time plot of log-scaled skull length illustrates the early expansion of ichthyosaur morphospace (Fig. 2). Across 108 phylogenies, the EHD model was overwhelmingly favored over Brownian Motion (BM), Ornstein-Uhlenbeck (OU), and Early Burst (EB) alternatives. EHD had the lowest AICc in 92% of cases when the best model was selected using ΔAICc = 0, and it was retained in 99% of cases when models within ΔAICc ≤ 2 were accepted. Mean AICc values further supported this result: EHD had the lowest mean AICc, 144.14, whereas EB was the next-closest model at 152.40, exceeding EHD by 8.26 AICc units and thus falling beyond the standard rejection threshold (Fig. 3B).

**Figure 2.**
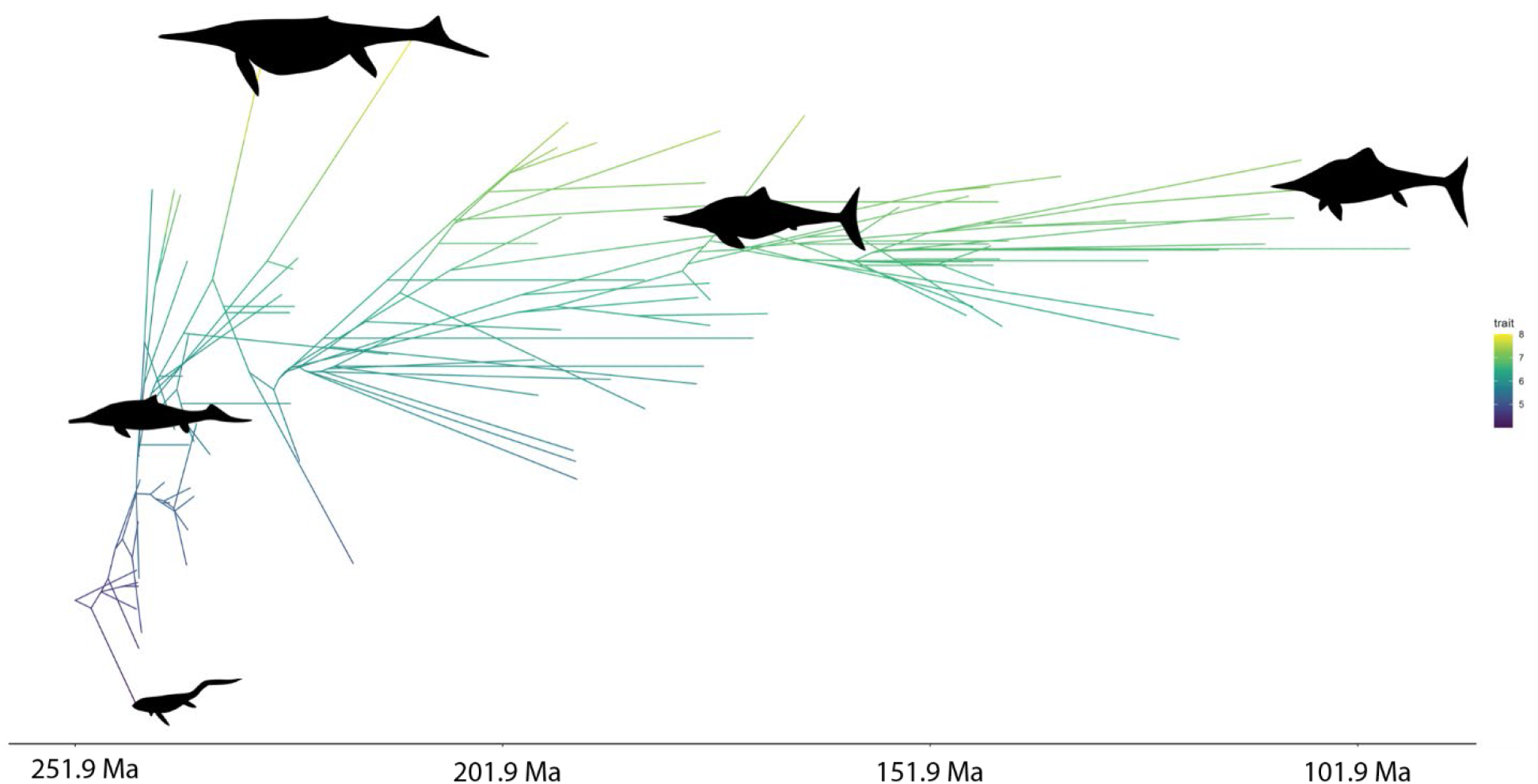
Phylomorphospace of skull length (log-transformed) in ichthyosaurs.

**Figure 3.**
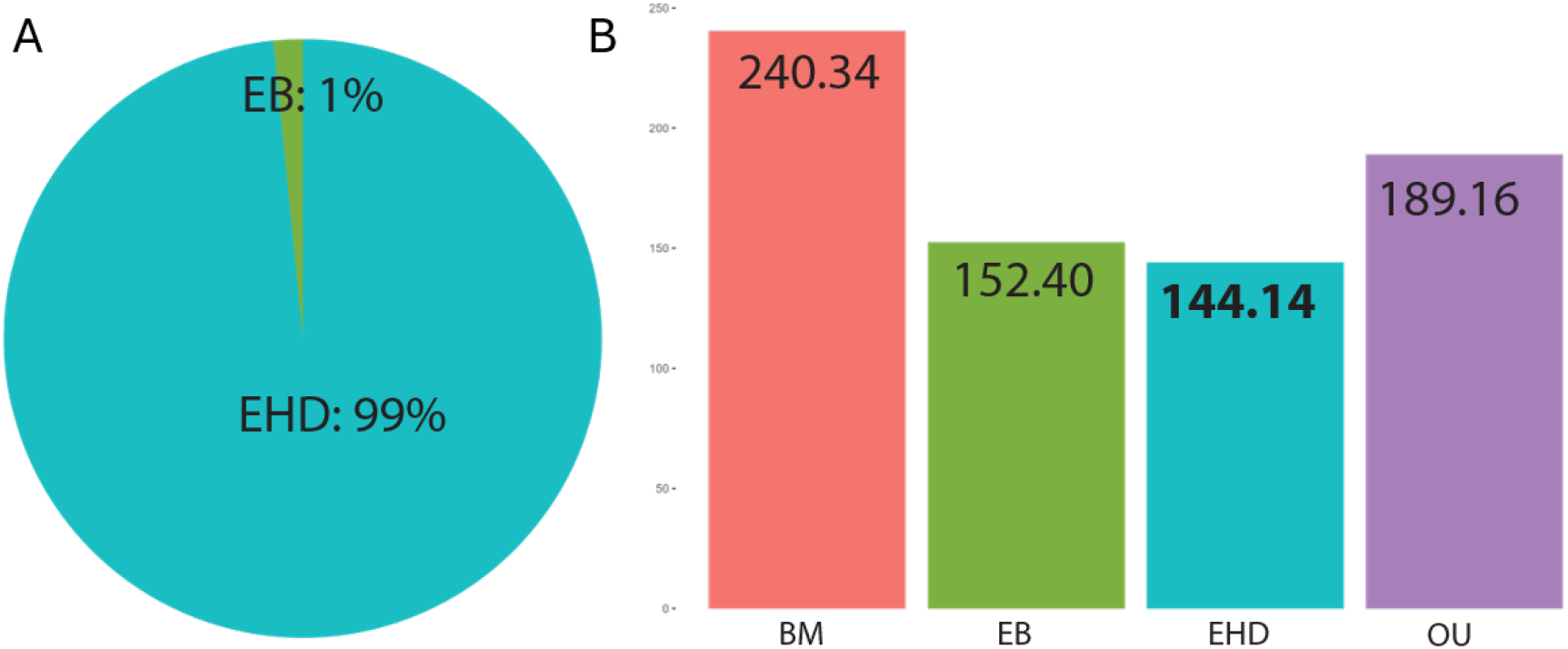
A.Pie chart of Akaike weights calculated from AIC_C_ values for each mode. On average, EHD is the best-fitting mode. B. AIC_C_ for each mode. BM: Brownian Motion; EB: Early Burst; EHD: Early High Disparity; OU: Ornstein Uhlenbeck.

We next estimated when disparity peaked under the EHD model. The median time of peak disparity, T_EHD_, was 231.1 Ma (Fig. 3A), calculated from the EHD peak-disparity equation using each model’s best-fit parameters. Across fitted EHD models, the average α and ρ values were 0.35 and 0.92, respectively (Fig. 4A). The estimated ancestral state was X0 = 4.34, while the optimum was θ = 7.07 (Fig. 4B). Because the median T_EHD_ occurs only ∼12.4% of the way through the total tree duration of 157.39 Myr, these results indicate a strongly bottom-heavy disparity profile, consistent with previous criteria for early high disparity (34, 35).

**Figure 4.**
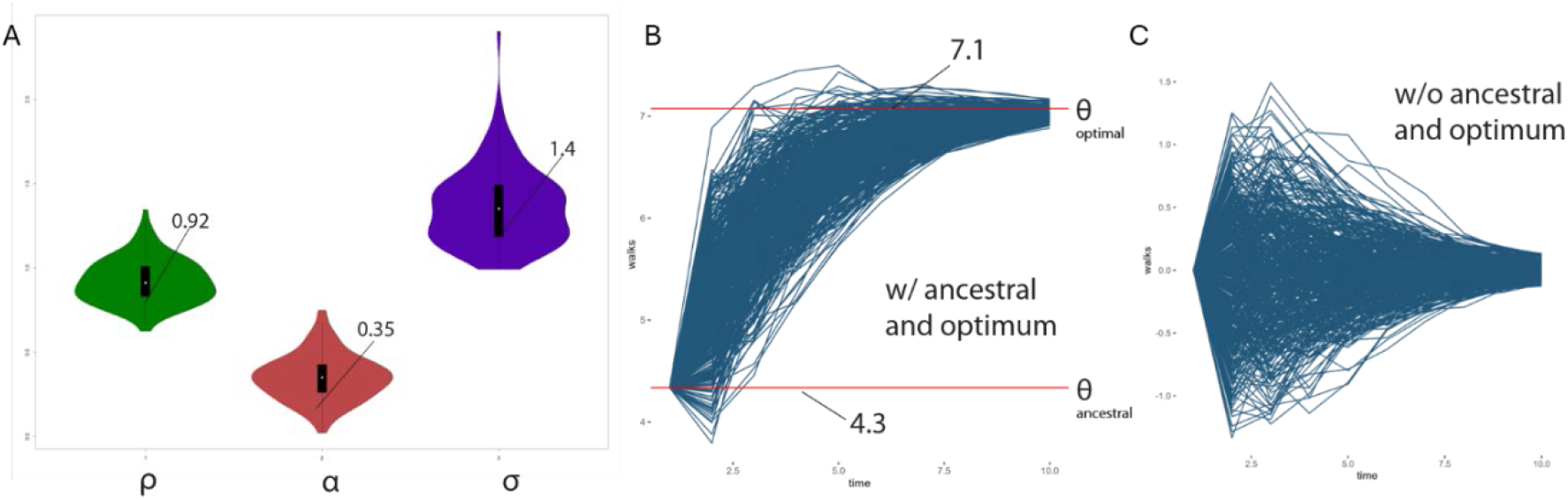
A.Best fitting EHD parameter distributions across 108 trees, excluding θ and X_0_. B. 500 Monte Carlo trait simulations of HD processes based on best fitting EHD parameters. θ_ancestral_: ancestral trait estimate; θ_optimal_: optimal trait estimate. C. Same as B, but excluding ancestral and optimal phenotypes in order to visualize the shape of the disparity profile.

Finally, we tested whether the estimated EHD peak coincided with major environmental or geological changes during the Triassic (Fig. 6). Changepoint analyses identified several shifts across the examined time series, but few aligned closely with T_EHD_ at 231.1 Ma. Sea level showed a single changepoint at 217.5 Ma (Fig. 6A), well after the estimated peak of disparity. A three-changepoint sea-level model, included to capture two troughs and one peak in the series, placed changepoints at 247.3 Ma, 238.5 Ma, and 219 Ma. Sea surface temperature showed a single changepoint at ∼237.2 Ma (Fig. 6B), with the three-changepoint model identifying additional shifts at 247 Ma, 239 Ma, and 206 Ma. The rock area showed a notable change point at 214 Ma (Fig. 6C), also substantially later than T_EHD_. Sequence boundary data, however, showed a significant shift at ∼237 Ma (Fig. 6D), close to the temperature change point and preceding the estimated EHD peak by approximately 6 Myr. Overall, these analyses suggest that although ichthyosaur disparity peaked early in the Triassic, this peak does not coincide precisely with most tested environmental or geological change points.

## Discussion

The strong support for the novel EHD model indicates that ichthyosaur disparity evolution was characterized by an early rise to high disparity, followed by a subsequent decline in both disparity and evolutionary rates. This pattern is also apparent qualitatively in the phylomorphospace-through-time plot, which shows a distinctly bottom-heavy disparity profile (Fig. 2). Together, these results support previous interpretations of ichthyosaurs as an explosive early radiation (3, 54, 55), while providing an explicit phylogenetic comparative framework for modeling that dynamic.

The estimated peak of disparity occurred in the Middle Triassic, consistent with recovery dynamics following the EPME and potentially linked to ecological restructuring around the Carnian Pluvial Episode. The median T_EHD_ falls within the first 12% of total ichthyosaur clade duration, providing strong evidence for a bottom-heavy profile under previous criteria for early high disparity (34, 35, 56–59). Although the average T_EHD_ estimate places maximum disparity during the Carnian stage of the Upper Triassic (237–227 Ma), some estimates approach the origin of ichthyosaurs (Fig. 4). Because T_EHD_ is jointly determined by α and ρ, parameter combinations that produce stronger early expansion and/or stronger subsequent constraint shift the inferred disparity peak closer to the Permian–Triassic boundary at 251.9 Ma. These earlier estimates may suggest a slight signal of an even more rapid initial expansion of ichthyosaur disparity immediately after the EPME.

If T_EHD_ estimates coincided with mass extinction boundaries, strong support for EHD could reflect extinction-driven shifts in macroevolutionary regime, such as a transition from EB-like expansion to OU-like constraint (60), rather than an intrinsic early-disparity dynamic. This is especially relevant because mass extinctions can reset macroevolutionary regimes broadly (61, 62). However, no T_EHD_ estimates approach the end-Triassic boundary (Fig. 3), indicating that the strong support for EHD is unlikely to be an artifact of the end-Triassic mass extinction.

This distinction is relevant given the ongoing debate over the effect of the end-Triassic mass extinction on ichthyosaur disparity. Some studies report a dramatic bottleneck in diversity and disparity across the Triassic–Jurassic boundary (2), whereas others find no significant decline in craniodental disparity across this interval (63). Our results are more consistent with the latter interpretation, in that the inferred disparity peak is not concentrated near the Triassic– Jurassic boundary. However, this comparison should be made with caution: previous studies primarily used discrete-character craniodental matrices to detect temporal shifts (63), whereas our analysis uses skull length. These differences suggest that distinct anatomical regions or trait types preserve different temporal signals of ichthyosaur disparity.

Although the timing of T_EHD_ do not correspond with the end-Permian or end-Triassic mass extinctions, ichthyosaur EHD may otherwise be influenced by abiotic, ecological, and evolutionary events occurring throughout the Triassic Period, such as the Mesozoic Marine Revolution (MMR) (31). The MMR encompassed a broad suite of ecological innovations that began during the Triassic. In marine vertebrates, these included fish-shaped body plans, high-velocity swimming, increased durophagy, and, in teleost fishes, protrusible upper jaws (20). Because the MMR is often interpreted as an escalating predator–prey “arms race” (19, 20), the early peak in ichthyosaur disparity may reflect rapid expansion into diverse foraging niches associated with increasingly mobile prey, including belemnites, ammonites, and fishes.

Subsequent declines in ichthyosaur disparity may have been linked to habitat restructuring and increasing ecological specialization. Across the Triassic, marine reptile ecomorphology shifted from durophagous and nearshore-associated forms toward more pelagic habits, a transition correlated with rapid sea-level fall and the reduction of shallow-water habitats for benthic feeding (30). More generally, sea-level regressions reduce habitat availability for nearshore taxa (64), potentially imposing selective pressures favoring open-ocean adaptations while disproportionately eliminating eel-like and lizard-like morphologies associated with shallower habitats in early ichthyosaurs (65). At the same time, the onset of the deep-marine “carbonate factory” (66), linked to the origin and expansion of dinoflagellates and coccolithophores (67), may have shifted productivity toward open-ocean systems. Under this scenario, ichthyosaur disparity decline from the Middle Triassic onward may reflect both extinction selectivity against nearshore morphologies and convergent evolution toward streamlined pelagic forms (68).

Thus, although our results do not support either the end-Permian or end-Triassic mass extinction as an artefactual source of EHD support, later Triassic events may still have influenced the inferred pattern. In particular, the Carnian Pluvial Episode could have contributed to the EHD signal if it increased extinction rates among taxa occupying the extremes of ichthyosaur disparity. Rather than contradicting the EHD interpretation, this possibility suggests that the observed pattern may reflect a combination of early ecological expansion, subsequent environmental restructuring, and selective extinction of morphologically extreme forms.

Time-series changepoint analyses of recent abiotic dataset (69) did not produce any clrar match to the disparity patterns in Ichthyosaurs. Sea level changepoints did not coincide directly with the mean T_EHD_ or its confidence interval (T_EHD_ CI: 230.75–232.2 Ma). However, the changepoint at 238.5 Ma precedes T_EHD_ and marks the transition from a rapid sea-level rise beginning after the 247 Ma lowstand to a subsequent interval of sea-level decline (Fig. 6A). This early Triassic transgression broadly parallels the initial rise in ichthyosaur disparity, even though the changepoint itself does not overlap the estimated peak.

This pattern suggests a plausible link between sea-level dynamics and the early expansion of ichthyosaur ecospace. Sea-level rise, or transgression, can increase shallow-marine and nearshore habitat area, thereby expanding ecological opportunity for diversification and disparification (70). Under this scenario, rising sea level from 247 Ma to 238.5 Ma may have promoted ichthyosaur diversification by increasing available nearshore ecospace. The subsequent sea-level fall from 238.5 Ma to 217 Ma may then have reduced nearshore habitats, favoring open-ocean adaptations while contributing to declining disparity among nearshore-associated forms.

This relationship appears to weaken after 217 Ma. From 217 Ma to the end of the Triassic, sea level rises sharply again, yet ichthyosaur disparity continues to decline or stabilize rather than recovering to earlier levels (Figs. 2, 5A). One possibility is that renewed nearshore ecospace preferentially benefited other Triassic marine reptile clades, such as eosauropterygians, including nothosaurs, placodonts, and early plesiosaurs. Alternatively, once ichthyosaurs became increasingly specialized for pelagic habitats, ecological overspecialization (71) may have reduced their evolutionary lability, limiting any subsequent return to nearshore adaptive zones.

**Figure 5.**
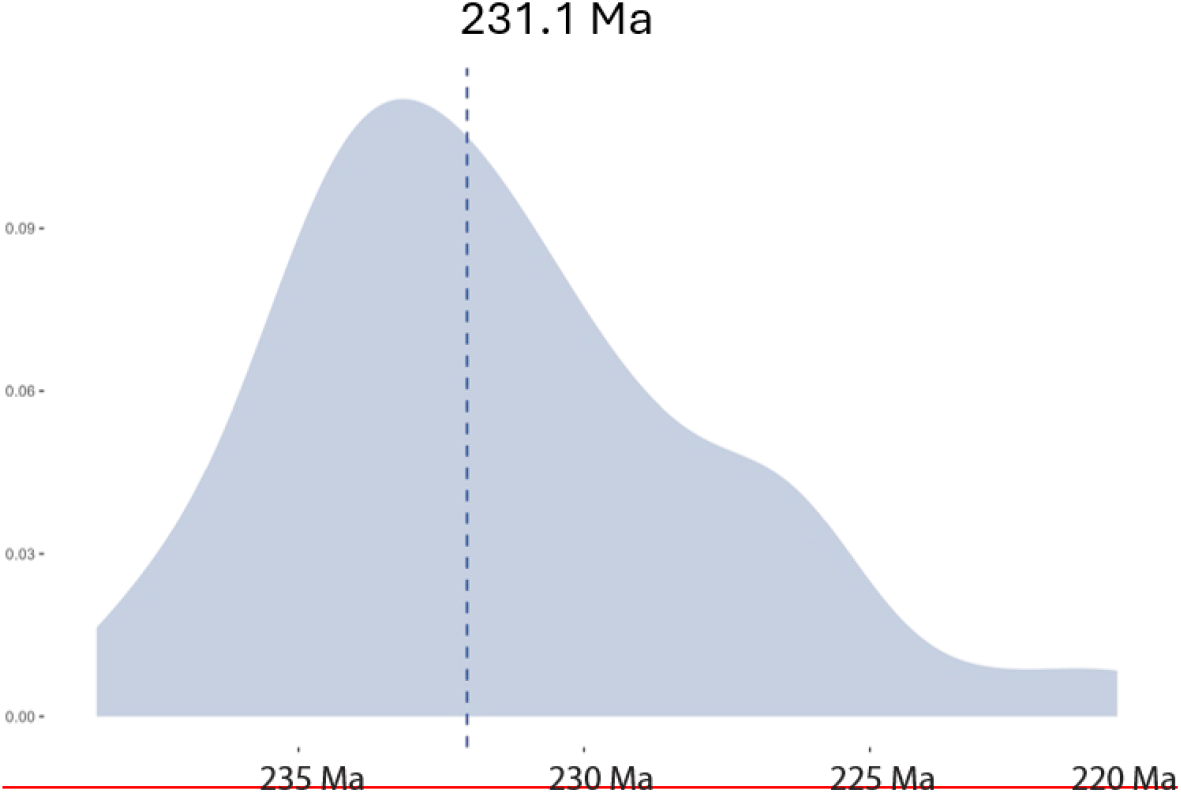
Distribution of T_EHD_, among 108 time-calibrated phylogenies. T_EHD_ represents the expected time of peak disparity, and is estimated from all best fitting parameter estimates of EHD.

**Figure 6.**
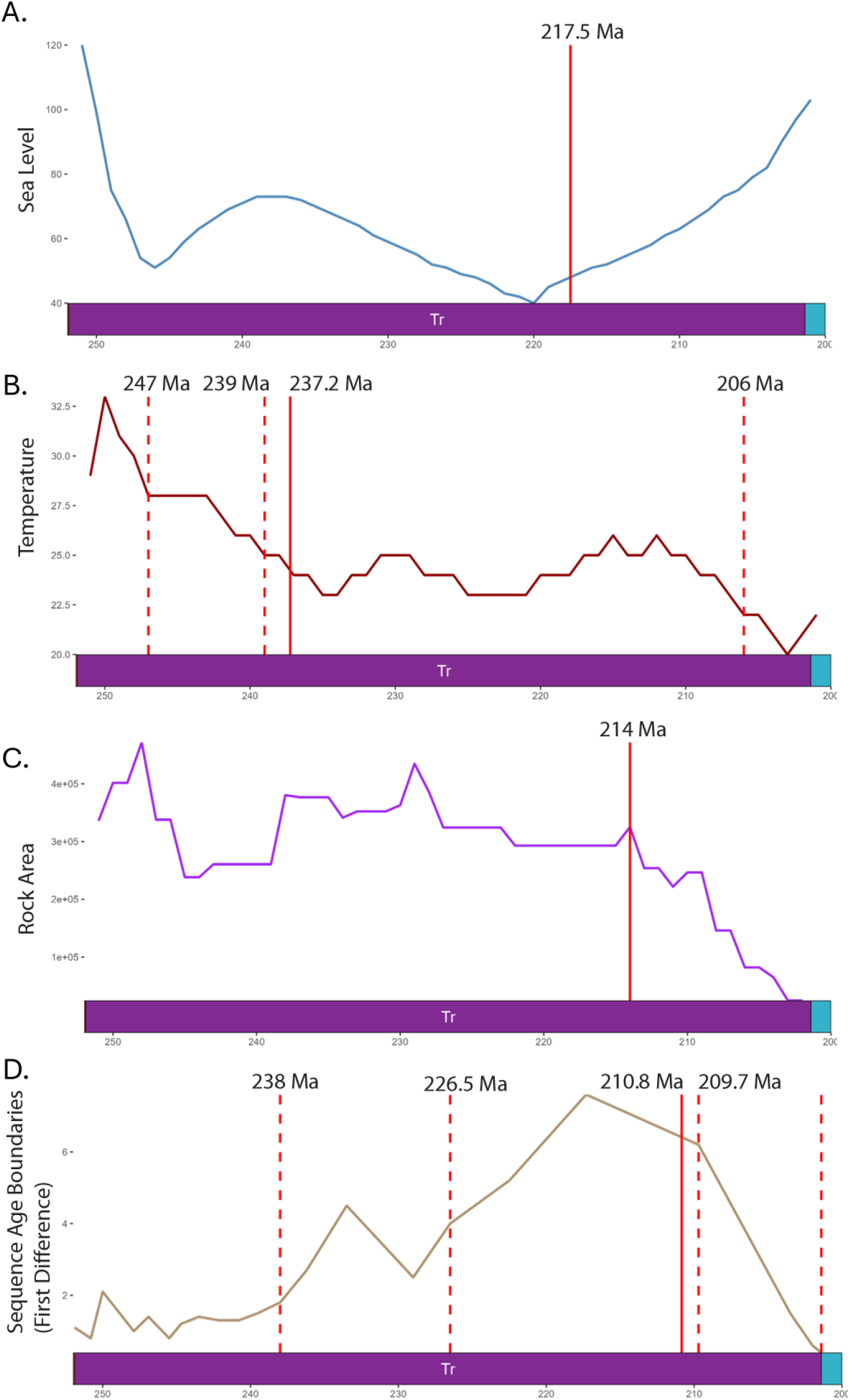
Time series with changepoints superimposed. A. Sea level throughout the Triassic Period. B. Sea surface temperature (red) throughout the Triassic, with one changepoint plotted (black dashed vertical). C. Marine rock area throughout the Triassic. D. Sequence age boundaries. Vertical solid lines: single significant changepoints detected. Vertical dashed lines: detected changepoints. Tr = Triassic Period.

The sea-surface temperature changepoint also precedes T_EHD_ but does not fall within the EHD peak-time confidence interval. Against a backdrop of global warming associated with Wrangellia terrane volcanism (23, 25), large-bodied ichthyosaurs, including shastasaurids and Cymbospondylus, may have experienced selection toward smaller body sizes. This would be consistent with Bergmann’s rule-like dynamics, in which smaller body sizes are favored in warmer environments and larger body sizes in cooler ones (72). Thus, although abiotic changepoints do not coincide precisely with T_EHD_, sea-level and temperature trends may still have shaped the ecological expansion and later constraint of ichthyosaur disparity during the Triassic.

In addition to abiotic time series for the Triassic, the potential role of geological biases in distorting the signal for true evolutionary modes was also assessed. Two primary time series were utilized: rock area and sequence age ranges (sequence “thickness”). Episodes of greater rock area availability are expected to contain greater samples of morphological varieties, just as greater taxic diversity is expected from larger rock area (32). The age range of marine sedimentary sequences may impact inferences of tempo and mode in phenotypic evolution by artefactually elevating evolutionary rates in times of sedimentary condensation or decreasing evolutionary rates in times of elevated sedimentary rates (33–35). The former scenario leads to incorrectly inferring evolutionary modes such as punctuated equilibrium (34), or perhaps rapid adaptive radiations similar to early bursts. The latter leads to incorrectly inferring episodes of stasis (33) and perhaps OU modes.

In the changepoint analyses of Triassic marine rock area, a significant changepoint occurs at 214 Ma. While a single significant changepoint in Triassic marine rock area occurs at 214 Ma, corresponding closely with a sea level changepoint (Fig. 6A,C), rock area shows a steady decrease while sea level simultaneously increases, contrary to intuitive expectations. This changepoint lies outside the confidence intervals for T_EHD_, suggesting an extremely low likelihood of strong impacts on disparity shifts. In fact, with the exception of the first ∼10 million years, rock area decreases before roughly stabilizing, then decreasing again for the remaining 10 million years of the Triassic (Fig. 6C). A decrease in rock area likely leads to a decrease in preservation of morphological varieties, whereas from the early to middle Triassic ichthyosaur disparity notably increases.

On the contrary, sequence age range experiences a changepoint at 238 Ma, much closer to the confidence interval of T_EHD_ (Fig. 6D). This corresponds not only to peak ichthyosaur disparity but also with the timing of the Carnian Pluvial Episode. The EHD model assumes exponentially elevated evolutionary rates prior to T_EHD_ then decreases in rates afterwards. Middle to Late Triassic sequences display considerably longer durations with fewer unconformities between major gaps (36). If Triassic sequences are the result of greater condensation and unconformity biases in the earlier portion of the Triassic, this may produce apparent EHD patterns. Alternatively, there may be a greater prevalence of deeper marine sequences in the latter half of the Triassic, consistent with sea level regression preventing sedimentary deposition in nearshore areas (37, 38). Nearshore and shallow oceanic areas experience greater fluctuations in sea level transgressions and regressions, or respectively deposition and erosion, fluctuations which produce greater incidences of uncomformities, while deeper water sedimentary sequences experience considerably less unconformities (39).

Geological biases and their impacts on disparity have been explored before, albeit not to the extent of fossil diversity. Previous works assessing the effects of geology on ichthyosaur phenotypic macroevolution have found an overall pattern of increasing then decreasing disparity throughout the Triassic remains after correcting for Lagerstätten effects (i.e. removing effects of geological formations with exceptional preservation) (40). Generally, disparity measures appear much less affected by geological sampling biases than diversity estimates (41), as morphological disparity largely concerns variance measures which have shown to be more robust towards sampling artefacts (42).

Ichthyosaurs, and other marine reptiles, naturally originated in nearshore environments as secondarily aquatic tetrapods, environments which are typically the focus of discerning common cause mechanisms from real biological or geological signal (43, 44). ‘Common cause’ theories posit that the same variables that drive sedimentary rock dynamics through time also drive diversification dynamics (45, 46). Invasion of new ecospaces, novel innovations, combined with the emptying of ecological niches due to mass extinctions, are factors considered primary influences on the tempo and mode of an adaptive radiation (22, 47). Ichthyosaurs potentially underwent tremendous disparification due to enhanced eco-evolutionary experimentation in nearshore environments, assuming sea level fluctuations lead to turnover through increased origination and extinction rates (38). Early Triassic ichthyosaurs tended to more lizard-like and eel-like swimming forms (48), but would also rapidly adapt to pelagic habitats (7), as typified by the parvipelvians with the stereotypical streamlined, fusiform morphology (more prominent later in the Triassic) (49). Dietarily, durophagous ichthyosaurs likely feeding on benthic marine invertebrates were far more common in the Early-Middle Triassic (12, 50), simultaneously ichthyosaurs were already starting to display increasingly cephalopod-reliant diets (51).

## Conclusions

An extensive macroevolutionary literature has developed around ichthyosaurs, including studies of evolutionary rates and modes (3, 4, 52), diversification (2, 53), ecomorphology and ecospace modelling (52, 54, 55) and geologic sampling biases (40, 56). The attention toward ichthyosaurs is due to their relatively complete fossil record (40), convergent evolution to modern marine vertebrates such as sharks, swordfish, and dolphins (57), and their explosive diversification shortly after the EPME (4). The estimated peak in disparity does not coincide with the end-Triassic mass extinction, suggesting that support for the EHD model is not simply an artifact of this event. Instead, the timing of T_EHD_ points to earlier Triassic dynamics, potentially including ecological and extinction-related changes associated with the Carnian Pluvial Episode (25).

An important next step is to evaluate how diversity dynamics and geological sampling interact with EHD model inference. Although our results do not indicate that these factors are driving the recovered EHD signal, they could still shape the broader pattern of disparity through time and influence model support. Previous studies have inferred early saturation of ichthyosaur diversification (34), and variation in species richness through time could affect the strength of support for an EHD model. If observed diversity patterns are partly shaped by geological sampling, then sampling structure could also influence apparent disparity dynamics. In this sense, the strong EHD signal recovered here provides a clear framework for future tests of how biological diversification, extinction selectivity, and fossil-record structure jointly shaped ichthyosaur disparity throughout the Triassic.

## Materials and Methods

### Comparative Data

Ichthyosaur skull lengths and time-calibrated phylogenies via the Hedman method (58) were extracted from (3). Each tree contains 108 icthyosaur taxa. Over half of ichthyosaur species (55)present in the phylogeny are missing skull length data. We subjected the entire dataset of trait data (skull length, total body length, orbit length, and orbital sclerotic diameter) to phylogenetic imputation via *Rphylopars* package (59) to increase the number of taxa included in model fitting.

### Early High Disparity Model

Our novel phylogenetic comparative model is an explicit combination of two stochastic differential equations (SDEs): Early Burst (27, 30) and Ornstein-Uhlenbeck (28, 29). EB processes describe exponentially decreasing evolutionary rate through time (-ρ), resulting in rapid increases of trait variance while maintaining a constant mean trait value (X_0_) (Fig. 1A) (27, 30). The expected mean trait in an OU process changes from an initial trait value (X_0_) towards an adaptively optimal value (θ) through a constraining force (α) which pulls the trait value towards θ while enforcing stochastically constant trait variance upon reaching θ (Fig. 1B) (28, 29, 60, 61). The new model presented here explicitly combines properties of these two modes, where the earliest portions of the process are dominated by EB-like dynamics, until reaching a peak trait variance and thereafter decreasing variance due to the constraining forces of the OU component of the model (Fig. 1C).

Below we present stochastic differential equations (SDEs) for OU, EB, and EHD processes, along with expected mean, variances and covariances for the EHD process. T represents the root-to-tip distance (age) of taxon *ii, CC*_*iiii*_ is the ‘raw’ phylogenetic covariance matrix, and *tt*_*iiii*_is the independent path length matrix. The derivation for T_EHD_ involves solving for time (T) when setting the derivative EHD variance equal to zero. See Appendix I for full derivation.

SDE (OU)

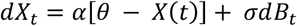

SDE (EB)

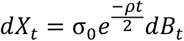

SDE (EHD)

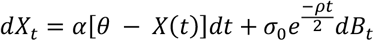

Expectation (Mean):

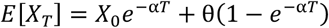

Variance:

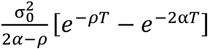

Covariance:

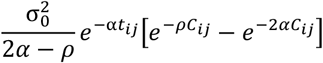

Expected Time of Peak Disparity:

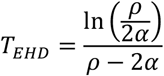

### Early High Disparity Model Fitting

We employ a maximum likelihood framework for estimating best-fitting parameters of the EHD mode and comparing relative model fit to BM, OU, and EB modes. The expected mean, variance, and covariance equations are typically used to calculate the phylogenetic covariance matrix (VCV) and expected means (62–64). This requires inverting of the phylogenetic VCV and calculating its determinant, the most computationally taxing steps in ML estimation (65–67).

Therefore, we employ an alternative approach using G_linv_ PCMs, a family of models specific to Gaussian-distributed traits, methods specific for evolutionary modes producing Gaussian (normal) distributions of traits. Each model must satisfy two conditions (66, 67):

1. Traits from two branching lineages are expected to evolve independently following branching from a node.
2. The mean and variance of a trait at any time *t* satisfy two further conditions:

a. The expectation (mean) is linearly dependent on the ancestral trait value.
b. The variance is independent of the ancestral trait.

Skull length data and time-calibrated phylogenies were subjected to maximum-likelihood based model-fitting and comparisons. The relative fit of BM, OU, EB, and EHD modes were evaluated using the R packages *PCMBase, PCMFit* (66, 67), and *PCMkappa* (68). G_linv_ implementations in *PCMBase* and *PCMFit* were designed with complex and parameter-rich modes in mind, as such they implement a tree traversal pruning algorithm assuming stochastic branching processes for greater computation ease. *PCMkappa* is a custom R package using *PCMBase* and *PCMFit* as dependencies, and includes custom functions for EB and EHD (69). Log-likelihoods were extracted to calculate AIC_c_ for each mode. ΔAIC_c_ was calculated by subtracting the minimum AIC_c_ from all competing model AIC_c_. The lowest AIC_c_ indicates the overall best-fitting mode, such that the mode with an ΔAIC_c_ = 0 indicates an overall best-fitting mode relative to all other modes fitted (70). Any modes with a ΔAIC_c_ > 2 are significantly worst fitting than the modes with ΔAIC_c_ = 0.

### Changepoint Analyses: Paleoenvironmental and Geological Time-Series

We used changepoint analyses to determine if timing of shifts in ichthyosaur EHD patterns (T_EHD_) coincide with shifts in timing of external paleoenvironmental or geological biases through time. Various changepoint methods exist (71), the most basic form performs automatic detection of significant changes in the slopes of linear regression curves (72). We applied single shift-point models for paleoenvironmental and geological time-series using the function *segmented* in the R package *segmented* (72). These time series included sea level curves and sea surface temperatures (73), rock area (74), and sedimentary sequence ages (36). Estimates of rock area were downloaded via Macrostrat (74). Triassic sedimentary sequence ages were extracted from (36), by taking first differences between sequence ages. All time series were truncated to the Triassic, where ichthyosaur disparity dynamics were most pronounced.

## Acknowledgments

R.C.E thanks David Polly for PhD supervision, PhD committee members Claudia Johnson, Jessica Miller-Camp, Matthew Hahn, Elizabeth Housworth, as well as Polly Lab members, Department of Earth & Atmospheric Sciences at Indiana University, Benjamin Moon and Thomas Stubbs for access to datasets and discussions.

## Data Availability

All code and data necessary to run analyses are available in the following link: https://doi.org/10.5281/zenodo.22896982.

