## Supplementary Information for "A novel phylogenetic comparative model reveals strong signals for early high disparity in ichthyosaurs"

#### **This PDF file includes:**

- Supporting text
- Appendix I
- Figures S1 to S6
- SI References

### Supporting Information Text

#### Details of the EHD Model

##### Brief Primer on BM, OU, and EB modes

Phylogenetic comparative methods (PCMs) and analyses are utilized to infer historical macroevolutionary processes such as phenotypic evolutionary rates and modes (1–3), estimation and correction for phylogenetic signal (4–6), evolution across adaptive landscapes (1, 7) diversification dynamics, such as extinction and speciation rates (8, 9) Among these, trait evolutionary modelling fit tip data to a phylogenetic tree in order to evaluate competing models which best explain trait evolution. The most common formulations of trait evolutionary modelling are represented primarily by three stochastic time-series models: Brownian Motion (2, 4, 5, 10) (O'Meara et al. 2006), Ornstein-Uhlenbeck (1, 7), and Early Burst (3, 6). The latter two models compose the EHD model presented here, which necessitates a brief discussion on basic properties of these modes (including BM), and their micro/macroevolutionary interpretations.

The primary evolutionary modes most frequently encountered in phylogenetic comparative methods are Brownian Motion, Ornstein-Uhlenbeck, and Early Burst. When fitting these modes to comparative datasets, it is important to understand that the expected moments derived directly from their respective stochastic differential equations act as descriptors of both the mean trait trajectory through time, but also most relevant, changes in expected trait variance through time. These stochastic processes are best understood by initially modelling them as stochastic univariate time-series, guided by their stochastic differential equation form. Brownian motion (Fig. 1) involves a stochastic time-series where each subsequent discrete time step is drawn from a normal distribution centered at zero and a variance: $dX_t \sim N(0, \sigma^2)$. This randomly drawn normal variate is added to the initial value of the random walk. Then another randomly drawn normal variate is added to this value. This process repeats a specified number of times, until a complete random walk is produced. This value is added to the last value in the process, continuing the 'random walk' (11).

$$dX_t = \sigma dB_t$$

An Ornstein-Uhlenbeck (OU) process, in which the trait value is pulled back toward an optimum value after an attempt to deviate from the optimum, can be viewed as an extension to Brownian motion random walks with the addition of special parameters such that the process tends toward the specified value (the optimum, $\theta$).

$$dX_t = \alpha[\theta - X(t)] + \sigma dB_t$$

The attraction towards $\theta$ is determined by $\alpha$, and causes an expected negative-exponential increase towards the optimum $\theta$. An OU process contains a random walk component ($\sigma dB_t$), which is added to a deterministic term containing $\alpha$ and $\theta$ terms (Eq. 2). The first step of an OU process is the same as in the random walk algorithm, where a random normal variate is drawn from a specified normal distribution (Fig. 2A.1). This variate is added to the previous value at $t - 1$, but the deterministic term of the OU process, $\alpha(\theta - X_t)$, is also added to these terms (Fig. 2A.1-3). This term contains the difference between the optimum $\theta$ and the value $X_t$ at time $t$ multiplied by an attraction parameter $\alpha$, which determines the rate at which the OU process moves towards $\theta$ and the variance of the process at time $t$ when reaching the optimum (1, 7). Therefore, there is an exponential increase in the OU process until reaching the optimal value, where the mean value stabilizes through time. Random deviation introduced by the Brownian motion component introduces fluctuations in the OU process away from the optimum, but is subsequently drawn back towards the optimum because of the attraction parameter $\alpha$ (hence its nickname, the "Rubber-Band Model").

The Early Burst process (3, 6) assumes the rate parameter, $\sigma^2$, decreases exponentially through time, such that:

$$\sigma^2(t) = \sigma_0^2 e^{-\rho t}$$

Each randomly drawn normally distributed variable is multiplied by $e^{-\rho t}$, and added to the previous value in the process (Fig. 3A). Much like in a Brownian motion/random walk, each time-series simulated randomly is not constrained to a particular trajectory. However, each simulated time-series approximates a negative exponential curve, with a rapid increase followed by a flattening of the curve. In SDE form, the early burst process takes the form:

$$dX_t = \sigma_0 e^{\frac{-\rho t}{2}} dB_t$$

Since Harmon et al. (2010), this equation has been the predominant formulation of the EB process. However, the equation is missing a crucial component of microevolutionary processes, which is stochastic noise around the respective asymptote of each EB walk. This may be illustrated in Fig. 3B, where complete flattening of each EB walk occurs. This peculiar behavior of EB walks is due to the effect of large $t$ on the exponential component in the EB equation: as $t \rightarrow -\infty$, $e^{-\rho t/2} \rightarrow 0$, subsequently removing the effect of $\sigma_0 dB_t$. This property is important to consider for maximum likelihood estimations and optimization algorithms, a property which likely results in optimization difficulty for converging on a reliable solution and substantially increasing the amount of time for MLE inference.

Monte Carlo simulations of each process inform us on the changes in variance of multiple stochastic time-series simulated simultaneously, and can help confirm expected means and variances at the end of these processes (Fig. 5). These SDEs also form the basis of calculating expectations (expected means), variances, and covariances through stochastic calculus (12). In phylogenetic comparative methods, theoretical expectations of these moments are incorporated into calculations of the phylogenetic covariance matrix (1, 13–15).

### APPENDIX I

#### Stochastic Calculus of the Early High Disparity Model

Presented below are the calculations for the expected mean, variance, and covariance of the early-high disparity model. This is composed of four primary steps: solving the SDE, then using this solution to calculate the expected mean, variance, and covariance utilizing stochastic calculus (12). These moments are used for implementation into the likelihood equation, given a time-calibrated phylogeny and univariate measurements per tip. This approach is currently implemented in various mode-fitting functions in R such as *fitContinuous()* in *geiger* (16) and *mvMORPH* functions such as *mvBM*, *mvOU*, *mvEB*, *mvSHIFT* (17), to list a few. Utilizing the phylogenetic covariance matrix is computationally taxing, however, as most mode-fitting functions necessitate inverting the phylogenetic covariance matrix and calculating the determinant of the phylogenetic covariance matrix (18). Several packages offer functions to substantially improve computational speed at no expense to reliable best-fitting parameter estimates, such as the “3-point covariance structure” approach (18), and the $G_{\text{linv}}$ family of Gaussian phylogenetic comparative methods, which use similar pruning algorithmic frameworks to estimating birth-death rates on a phylogeny (19). The latter method is actually used in this work, as opposed to the “traditional” approaches which can be found packages such as *geiger* (16), *hansen* (1), and *OUwie* (20).

Although one does not need explicit phylogenetic covariance formulas for the newer methods mentioned above, the calculations for the expectation and variance will prove useful for calculating expected times of maximum disparity of the EHD mode ($T_{\text{EHD}}$). This parameter is derived by setting the derivative of the EHD variance formula to zero, then solving for $t$ (Appendix I). For convenience, I will summarize these equations below, including the EHD covariance formulas.

$\rho$

SDE Solution:

$$X_T = e^{-\alpha T} X_0 + \theta(1 - e^{-\alpha T}) + \sigma_0 \int_0^T e^{(\alpha - \frac{\rho}{2})t} e^{-\alpha t} dB_t$$

Expectation (Mean):

$$E[X_T] = X_0 e^{-\alpha T} + \theta(1 - e^{-\alpha T})$$

Variance:

$$\frac{\sigma_0^2}{2\alpha - \rho} [e^{-\rho T} - e^{-2\alpha T}]$$

Covariance:

$$\frac{\sigma_0^2}{2\alpha - \rho} e^{-\alpha t_{ij}} [e^{-\rho C_{ij}} - e^{-2\alpha C_{ij}}]$$

Where T represents the root-to-tip distance (age) of taxon $i$, $C_{ij}$ is the 'raw' phylogenetic covariance matrix, and $t_{ij}$ is the independent path length matrix. Explanations are presented in Figs. \_ and \_, and in the text.

**Solution to SDE:**

$$dX_t = \alpha[\theta - X(t)]dt + \sigma_0 e^{-\rho t/2} dB_t$$

$$dX_t = [\alpha\theta - \alpha X(t)]dt + \sigma_0 e^{-\frac{\rho t}{2}} dB_t$$

$$dX_t = \alpha\theta dt - \alpha X(t)dt + \sigma_0 e^{-\frac{\rho t}{2}} dB_t$$

$$dX_t + \alpha X(t)dt = \alpha\theta dt + \sigma_0 e^{-\frac{\rho t}{2}} dB_t$$

$$e^{\alpha t}(dX_t + \alpha X(t)dt) = e^{\alpha t}(\alpha\theta dt + \sigma_0 e^{-\frac{\rho t}{2}} dB_t)$$

$$d(e^{\alpha t} X(t)) = e^{\alpha t} \alpha\theta dt + \sigma_0 e^{\alpha t} e^{-\frac{\rho t}{2}} dB_t$$

$$\int_0^T d(e^{\alpha t}) = \int_0^T e^{\alpha t} \alpha\theta dt + \int_0^T \sigma_0 e^{\alpha t} e^{-\frac{\rho t}{2}} dB_t$$

$$e^{\alpha T} X_T - e^0 X_0 = \alpha\theta \frac{e^{\alpha T} - 1}{\alpha} + \sigma_0 \int_0^T e^{(\alpha - \frac{\rho}{2})t} dB_t$$

$$e^{\alpha T} X_T - X_0 = \theta(e^{\alpha T} - 1) + \sigma_0 \int_0^T e^{(\alpha - \frac{\rho}{2})t} dB_t$$

$$e^{\alpha T} X_T - X_0 = \theta(e^{\alpha T} - 1) + \sigma_0 \int_0^T e^{(\alpha - \frac{\rho}{2})t} dB_t$$

$$e^{\alpha T} X_T = X_0 + \theta(e^{\alpha T} - 1) + \sigma_0 \int_0^T e^{(\alpha - \frac{\rho}{2})t} dB_t$$

$$X_T = e^{-\alpha T} X_0 + \theta(1 - e^{-\alpha T}) + \sigma_0 \int_0^T e^{(\alpha - \frac{\rho}{2})t} e^{-\alpha t} dB_t$$

**Expected Mean**

$$E[X_T] = X_0 e^{-\alpha T} + \theta(1 - e^{-\alpha T})$$

**Variance**

$$\text{Var}[X_T] = E[(X_T - E[X_T])^2]$$

$$= E\left[\left(\sigma_0 \int_0^T e^{(\alpha - \frac{\rho}{2})t} e^{-\alpha t} dB_t\right)^2\right]$$

$$= \sigma_0^2 e^{-2\alpha T} \int_0^T e^{2(\alpha - \frac{\rho}{2})t} dt$$

$$\begin{aligned} &= \sigma_0^2 e^{-2\alpha T} \left[ \frac{e^{(2\alpha-\rho)T} - e^0}{2\alpha - \rho} \right] \\ &= \frac{\sigma_0^2}{2\alpha - \rho} [e^{-2\alpha T} e^{(2\alpha-\rho)T} - e^{-2\alpha T}] \\ &= \frac{\sigma_0^2}{2\alpha - \rho} [e^{-\rho T} - e^{-2\alpha T}] \end{aligned}$$

**Covariance**

$$\begin{aligned} \text{Cov}[\mathbf{X}_T] &= \sigma_0^2 E \left[ \left( \int_0^i e^{(\alpha-\frac{\rho}{2})u-\alpha i} dB_u \right) \left( \int_0^j e^{(\alpha-\frac{\rho}{2})u-\alpha j} dB_u \right) \right] \\ &= \sigma_0^2 e^{-2\alpha i} e^{-2\alpha j} E \left[ \left( \int_0^i e^{(\alpha-\frac{\rho}{2})u} dB_u \right) \left( \int_0^j e^{(\alpha-\frac{\rho}{2})u} dB_u \right) \right] \\ &= \sigma_0^2 e^{-\alpha(i+j)} E \left[ \left( \int_0^{C_{ij}} e^{(\alpha-\frac{\rho}{2})u} dB_u + \int_{C_{ij}}^i e^{(\alpha-\frac{\rho}{2})u} dB_u \right) \left( \int_0^{C_{ij}} e^{(\alpha-\frac{\rho}{2})u} dB_u + \int_{C_{ij}}^j e^{(\alpha-\frac{\rho}{2})u} dB_u \right) \right] \\ &= \sigma_0^2 e^{-\alpha(i+j)} E \left[ \left( \int_0^{C_{ij}} e^{(\alpha-\frac{\rho}{2})u} dB_u \right)^2 + \left( \int_0^{C_{ij}} e^{(\alpha-\frac{\rho}{2})u} dB_u \right) \left( \int_{C_{ij}}^j e^{(\alpha-\frac{\rho}{2})u} dB_u \right) \right. \\ &\quad \left. + \left( \int_0^{C_{ij}} e^{(\alpha-\frac{\rho}{2})u} dB_u \right) \left( \int_{C_{ij}}^i e^{(\alpha-\frac{\rho}{2})u} dB_u \right) + \left( \int_{C_{ij}}^j e^{(\alpha-\frac{\rho}{2})u} dB_u \right) \left( \int_{C_{ij}}^i e^{(\alpha-\frac{\rho}{2})u} dB_u \right) \right] \\ &= \sigma_0^2 e^{-\alpha(i+j)} E \left[ \left( \int_0^{C_{ij}} e^{(\alpha-\frac{\rho}{2})u} dB_u \right)^2 \right] \\ &= \sigma_0^2 e^{-\alpha(i+j)} E \left[ \int_0^{C_{ij}} e^{2(\alpha-\frac{\rho}{2})u} du \right] \\ &= \frac{\sigma_0^2 e^{-\alpha(t_i+t_j+2C_{ij})}}{2\alpha - \rho} [e^{(2\alpha-\rho)C_{ij}} - 1] \\ &= \frac{\sigma_0^2 e^{-\alpha(t_{ij}+2C_{ij})}}{2\alpha - \rho} [e^{(2\alpha-\rho)C_{ij}} - 1] \\ &= \frac{\sigma_0^2}{2\alpha - \rho} e^{-\alpha t_{ij}} e^{-2\alpha C_{ij}} [e^{(2\alpha-\rho)C_{ij}} - 1] \\ &= \frac{\sigma_0^2}{2\alpha - \rho} e^{-\alpha t_{ij}} [e^{-\rho C_{ij}} - e^{-2\alpha C_{ij}}] \end{aligned}$$

**Early High Disparity Phylogenetic Covariance Matrix**

$$V = \frac{\sigma_0^2}{2\alpha - \rho} \begin{bmatrix} [e^{-\rho t_{ii}} - e^{-2\alpha t_{ii}}] & e^{-\alpha t_{ij}} [e^{-\rho C_{ij}} - e^{-2\alpha C_{ij}}] \\ e^{-\alpha t_{ij}} [e^{-\rho C_{ij}} - e^{-2\alpha C_{ij}}] & [e^{-\rho t_{jj}} - e^{-2\alpha t_{jj}}] \end{bmatrix}$$

**Derivation of $T_{\text{EHD}}$**

**Part 1: Calculate derivative for EHD Variance**

$$\begin{aligned} \text{Var}[\mathbf{X}_T] &= f(T) = \frac{\sigma_0^2}{2\alpha - \rho} [e^{-\rho T} - e^{-2\alpha T}] \\ \frac{df}{dT} &= \frac{\sigma_0^2}{2\alpha - \rho} \cdot \frac{d}{dT} [e^{-\rho T} - e^{-2\alpha T}] \\ \frac{d}{dT} [e^{-\rho T} - e^{-2\alpha T}] &= -\rho e^{-\rho T} + 2\alpha e^{-2\alpha T} \\ \frac{df}{dT} &= \frac{\sigma_0^2}{2\alpha - \rho} [2\alpha e^{-2\alpha T} - \rho e^{-\rho T}] \end{aligned}$$

Part 2: Set derivative equal to zero and solve for T.

$$\frac{\sigma_0^2}{2\alpha - \rho} [2\alpha e^{-2\alpha T} - \rho e^{-\rho T}] = 0$$

$$2\alpha e^{-2\alpha T} - \rho e^{-\rho T} = 0$$

$$2\alpha e^{-2\alpha T} = \rho e^{-\rho T}$$

$$2\alpha = \rho e^{(\rho - 2\alpha)T}$$

$$\frac{\rho}{2\alpha} = e^{(\rho - 2\alpha)T}$$

$$(\rho - 2\alpha)T = \ln\left(\frac{\rho}{2\alpha}\right)$$

$$T_{EHD} = \frac{\ln\left(\frac{\rho}{2\alpha}\right)}{\rho - 2\alpha}$$

### Figures

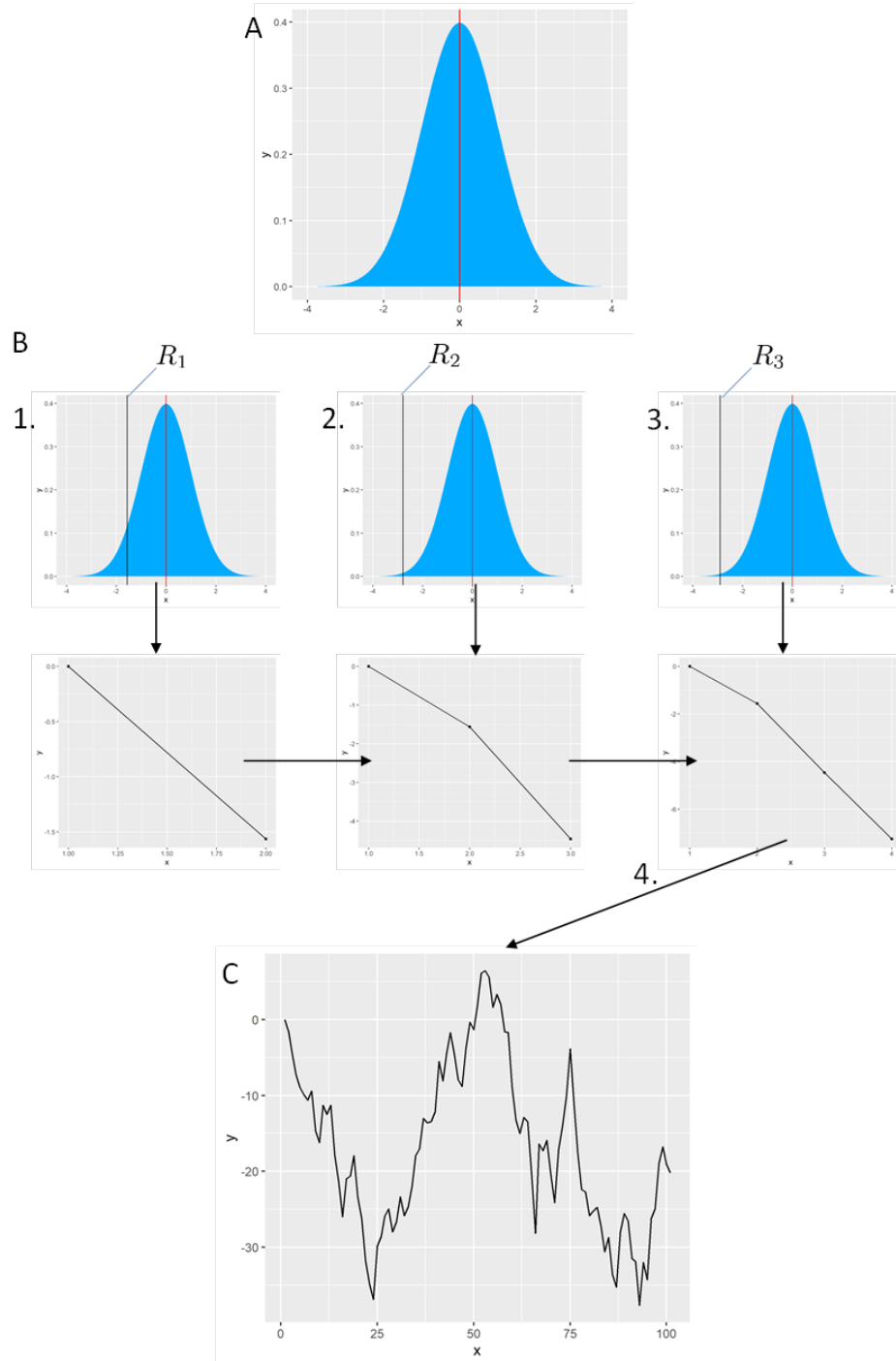

**Fig. S1.** Building a random walk (single Brownian motion).

**A.** A normal distribution with $\mu = 0$, and a user-defined $\sigma^2$. **B.** To build the random walk, random normal variates ($R_t$) are drawn from the normal distribution in A. 1. The first random normal variate ($R_1$) is drawn and added to the initial value of the random walk (in this case, $X_0 = 0$). In other words, $X_1 = X_0 + R_1$. 2. $R_2$ is drawn and added to the previous value at $t = 1$; $X_2 = X_1 + R_2$. 3. As in 2, $R_3$ is added to the previous value in the random walk ($X_3 = X_2 + R_3$). **C.** These steps are iterated several times (in this case, 100 steps), resulting in the random walk seen here.

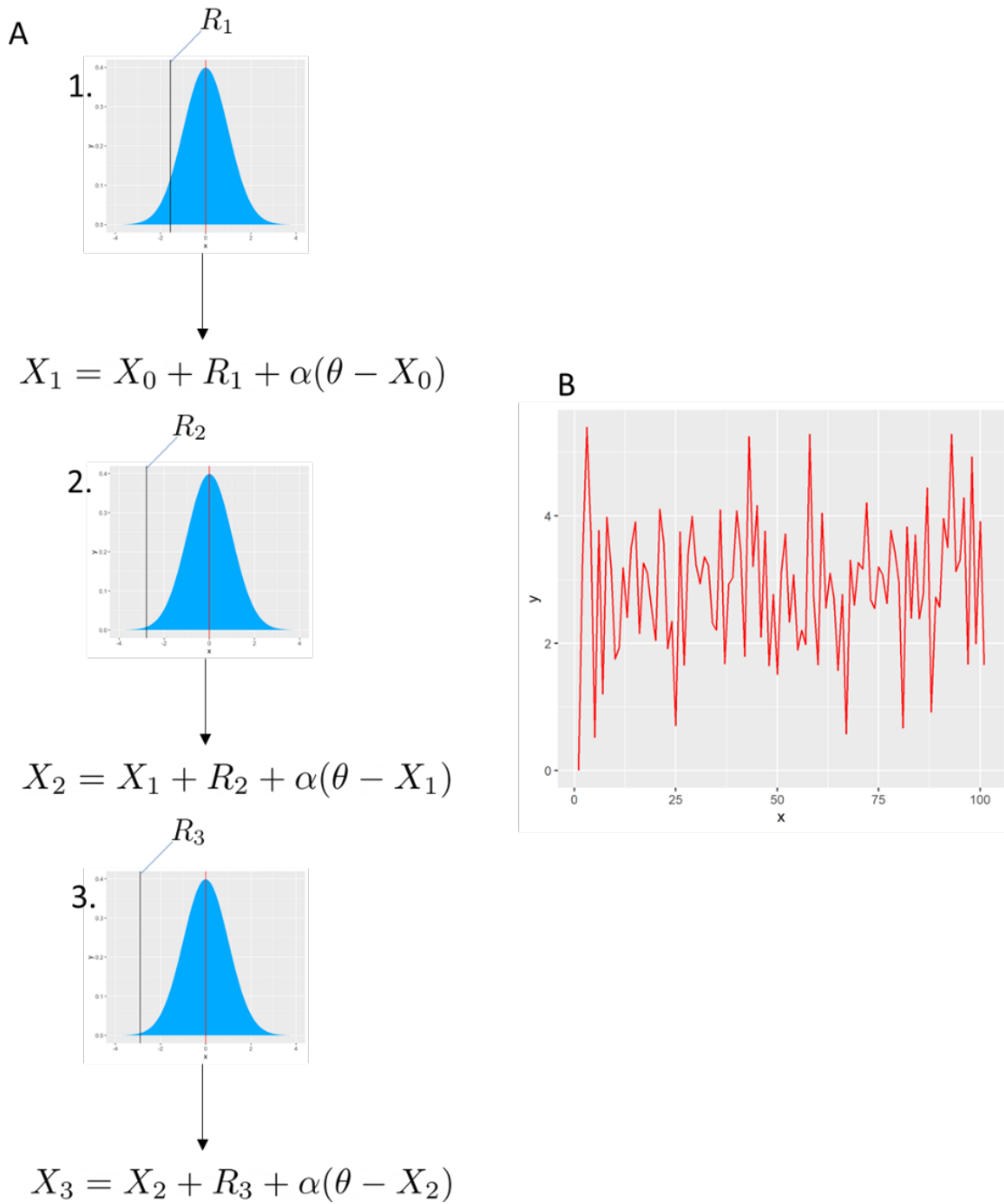

**Fig. S2.** Building an Ornstein-Uhlenbeck (OU) walk.

**A.** As in Fig. 1, a normal distribution is defined ($\sim N(\mu = 0, \sigma^2)$) from which random normal variates will be drawn. At each step however, the OU parameters are added to this normal variate.

1. $X_1 = X_0 + R_1 + \alpha(\theta - X_0)$.

2. $X_2 = X_1 + R_2 + \alpha(\theta - X_1)$

3. $X_3 = X_2 + R_3 + \alpha(\theta - X_2)$

**B.** The process outlined in A, but with a total of 100 steps.

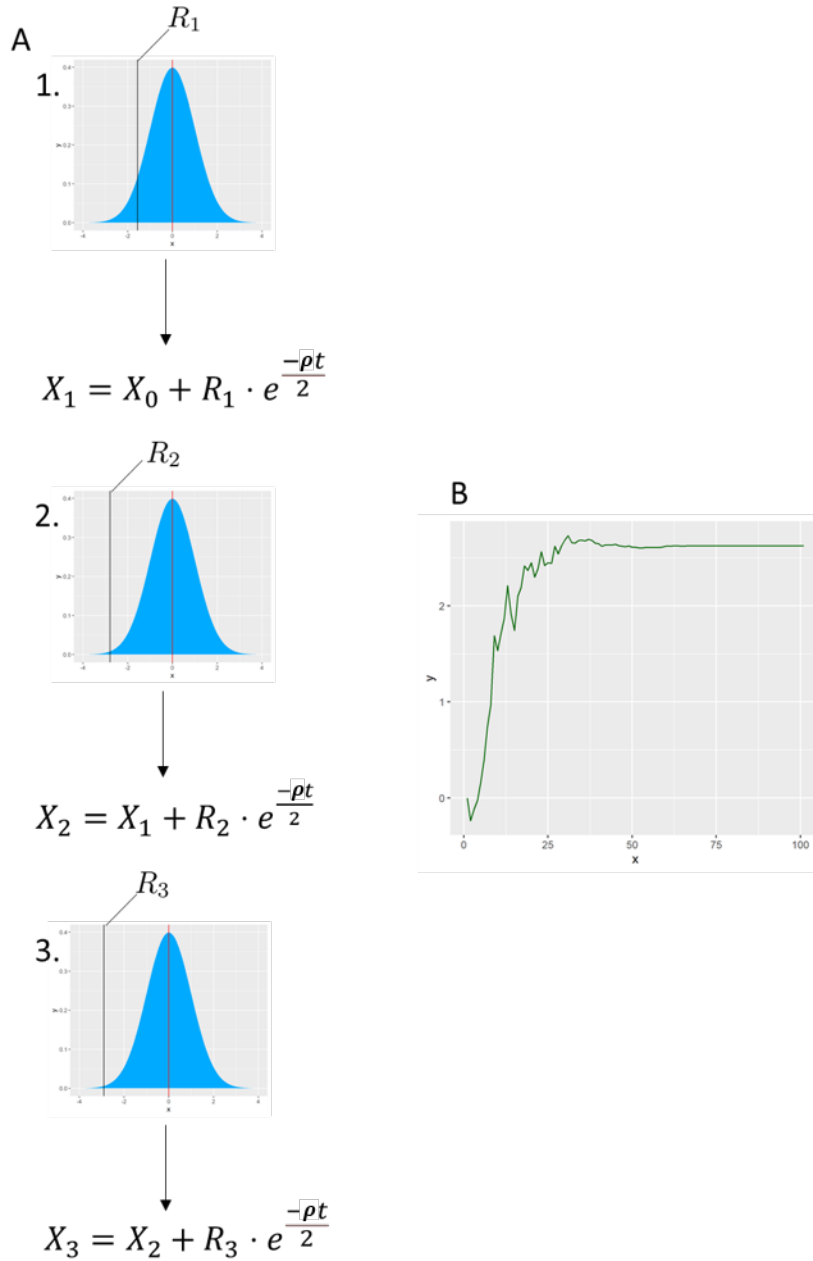

**Fig. S3.** Building an Early Burst (EB) walk.

**A.** As in Figs. 1-2, A random normal variate is drawn from a defined normal distribution ($\sim N(\mu = 0, \sigma^2)$). At each step, the normal variate is multiplied by the EB parameters.

1. $X_1 = X_0 + R_1 \cdot e^{\frac{-\rho t}{2}}$

2. $X_2 = X_1 + R_2 \cdot e^{\frac{-\rho t}{2}}$

3. $X_3 = X_2 + R_3 \cdot e^{\frac{-\rho t}{2}}$

**B.** The process outlined in A, but with a total of 100 steps.

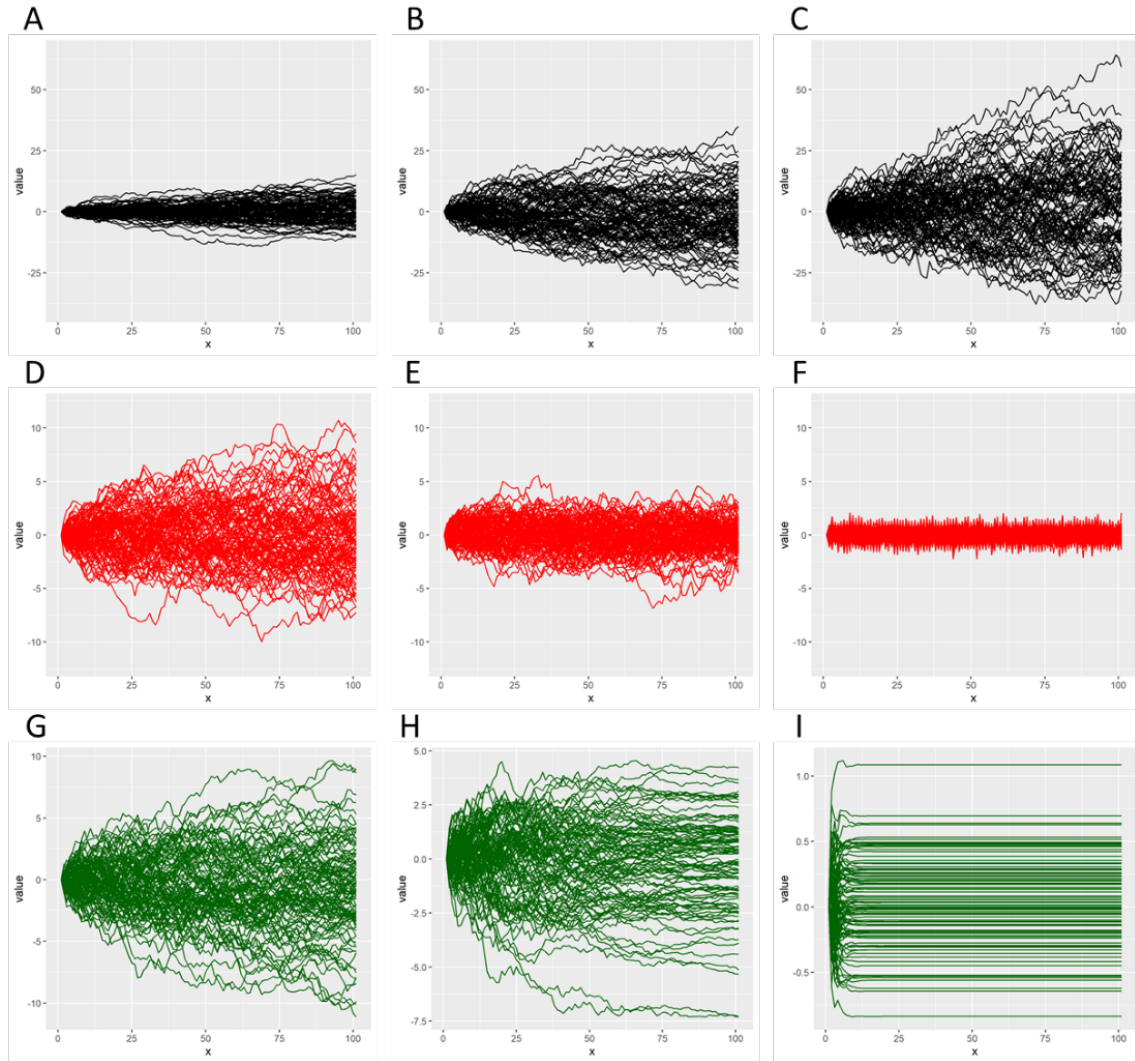

**Fig. S4. Monte Carlo univariate trait simulations for three canonical modes of evolution frequently applied in phylogenetic comparative studies.**

**A-C: Brownian Motion Monte Carlo simulations ($n = 100$).** A. $\sigma^2 = 0.25$. B. $\sigma^2 = 1.5625$. C. $\sigma^2 = 2$. The effects of increasing the variance of the normal distribution from which random normal variates are drawn is increasing the variance of the simulated random walks at each time step.

**D-F: Ornstein-Uhlenbeck Monte Carlo simulations ($n = 100$, $\sigma^2 = 0.25$ for Figs. D-F).** D. $\alpha = 0.01$. E. $\alpha = 0.05$. F. $\alpha = 1.5$. The effects of increasing the alpha parameter is to decrease the variance of the OU walks at each time step, and importantly constrains the variance for the duration of the OU walk, once all processes have reached the optimum $\theta$.

**G-I: Early Burst Monte Carlo simulations ($n = 100$, $\sigma^2 = 0.25$).** G. $\rho = -0.01$. H. $\rho = -0.05$. I. $\rho = -1$. As a result of decreasing $\rho$, all EB walks assume greater negative exponential profiles and reach their respective asymptotes more rapidly.

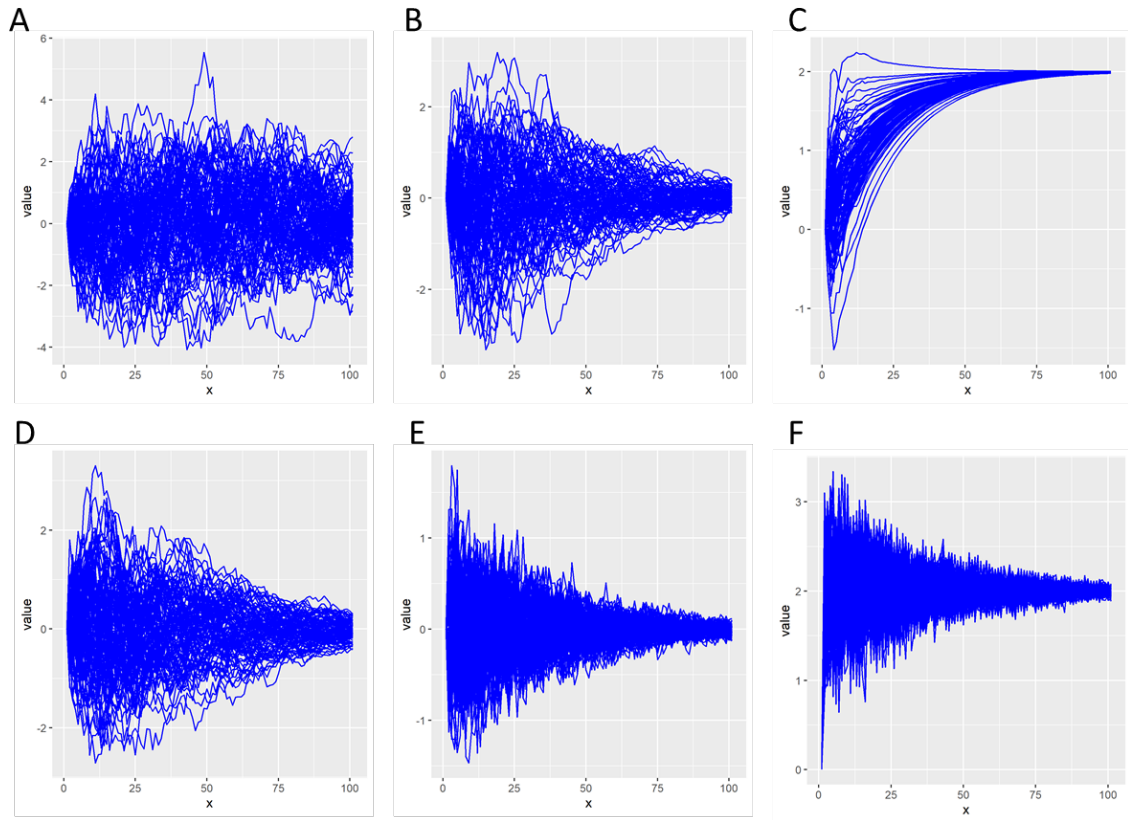

**Fig. S5.** Early High Disparity Monte Carlo simulations ($n = 100$, $\sigma^2=0.25$). **A.** $\alpha = 0.05$, $\rho = -0.01$. **B.** $\alpha = 0.05$, $\rho = -0.05$. **C.** $\alpha = 0.05$, $\rho = -0.5$, $\theta = 2$. **D.** $\alpha = 0.05$, $\rho = -0.05$. **E.** $\alpha = 0.5$, $\rho = -0.05$. **F.** $\alpha = 1$, $\rho = -0.05$, $\theta = 2$. A-C show results for varying only $r$, D-F show results for varying $\alpha$, and C, F have an optimum at $\theta = 2$.

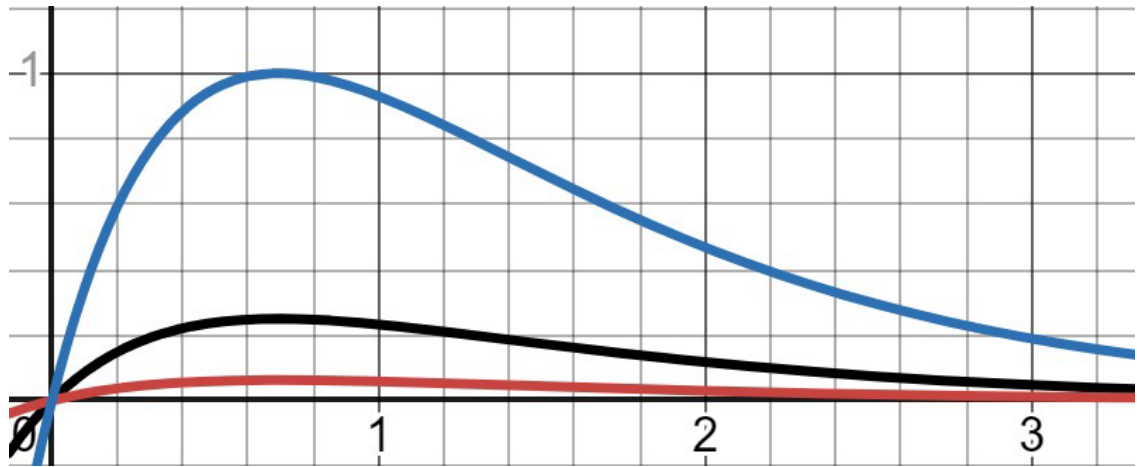

**Fig. S6.** Expected variance-through-time of EHD processes. Showcases the effect of varying $\sigma^2$ on the peak of maximum disparity. Black line: $\sigma^2 = 1$. Red line: $\sigma^2 = 0.5$. Blue line: $\sigma^2 = 2$.
